# A generalized kinetic model describes ion-permeation mechanisms in various ion channels

**DOI:** 10.1101/2021.05.08.443239

**Authors:** Di Wu

## Abstract

Ion channels conduct various ions across biological membranes to maintain the membrane potential, to transmit the electrical signals, and to elicit the subsequent cellular responses by the signaling ions. Ion channels differ in their capabilities to select and conduct ions, which can be studied by the patch-clamp recording method that compares the current traces responding to the test voltage elicited at different conditions. In these experiments, the current-voltage curves are usually fitted by a sigmoidal function containing the Boltzmann factor. This equation is quite successful in fitting the experimental data in many cases, but it also fails in several others. Regretfully, some useful information may be lost in these data, which otherwise can reveal the ion-permeation mechanisms. Here we present a generalized kinetic model that captures the essential features of the current-voltage relations and describes the simple mechanism of the ion permeation through different ion channels. We demonstrate that this model is capable to fit various types of the patch-clamp data and explain their ion-permeation mechanisms.

## Introduction

Cell membranes, made of lipid bilayers, are impermeable to inorganic ions. Various ions cross the membrane via the specific ion channels down their electrical-chemical gradient, or against this gradient at the expense of the extra energy, e.g., via the hydrolysis of the ATP molecules. Due to the availabilities and functions of different ion channels, pumps, and transporters at different locations, the concentrations of the various ions are maintained at the different levels in specific locations separated by the cell membranes, making ions as the important signaling molecules. In these processes, the transmembrane ion channels play important roles, owning to their varied capabilities to select and conduct ions across the membrane. The dysfunction of the ion channels can lead to various diseases (Ackerman & Clapham, 1997; Ashcroft, 2006; Lehmann-Horn & Jurkat-Rott, 1999).

Ion channel functions are usually studied by the patch-clamp recording method (Neher & Sakmann, 1976; Neher, Sakmann, & Steinbach, 1978), where the electrical currents are recorded responding to a series of the test voltages elicited. The shapes of the current-voltage curves (typical curves are shown in Fig. 1) are helpful in elucidating the channel functions, e.g., they can provide the information such as the magnitude of the inward and outward currents indicating the inward or outward rectifications, the steepness of the current slopes, the reversal voltage, etc. To compare the functions of the wild-type and mutant channels, or to compare the same channel under the different situations, multiple curves are usually plotted together for a quick and qualitative description of the channel functions. For the quantitative comparisons, and for the explanation of the ion-permeation mechanisms, we must use a model that fits the data.

**Figure 1.**
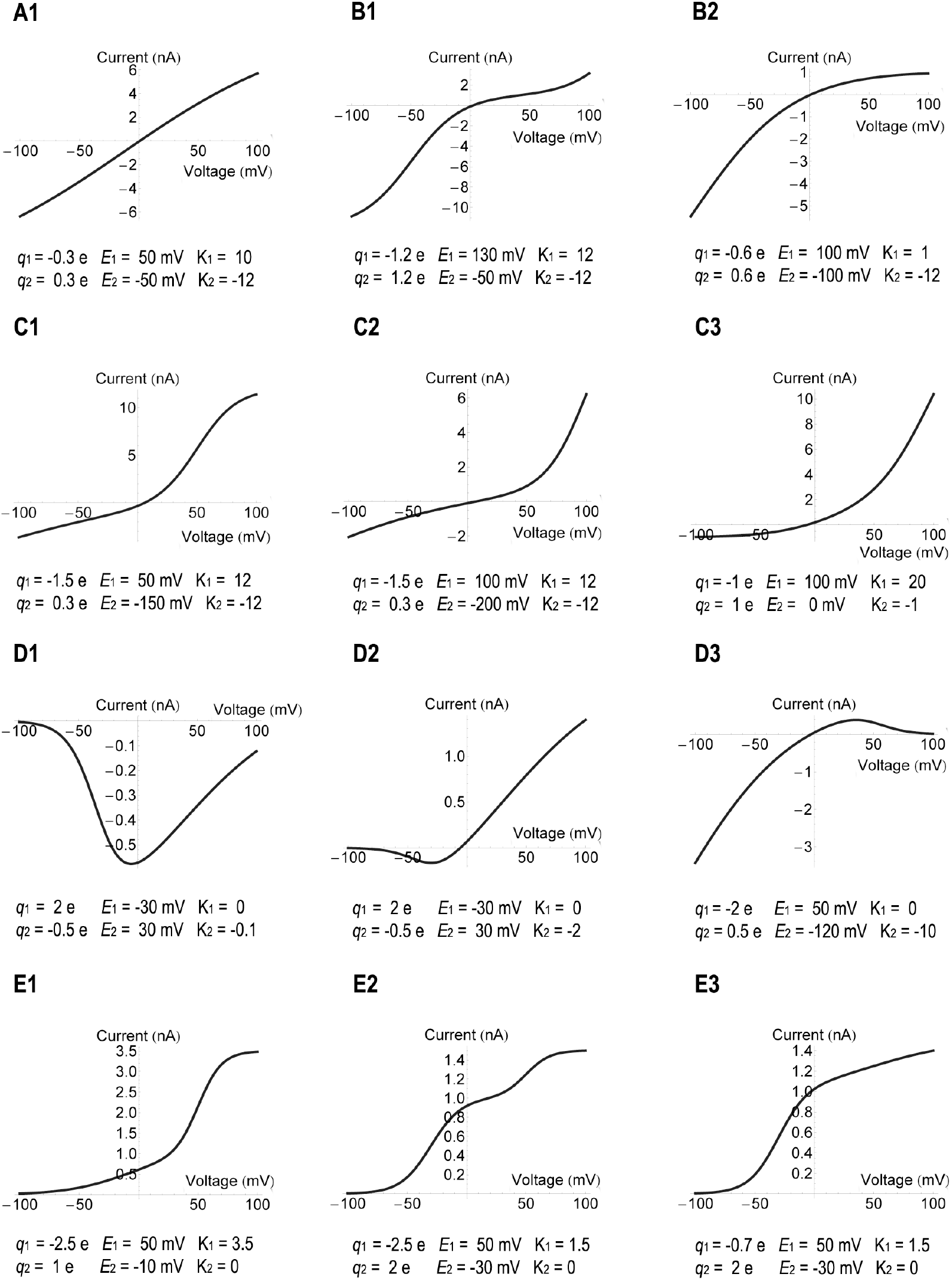
Typical current-voltage curves represented by the three-state model. Mechanism m3A3 was used for curves D1 and D2, and mechanism m3A2 was used for all other curves. The model parameters are listed below each curve. The current has the same unit as Ea, which was set arbitrarily to 1 nA for mechanism m3A2 and -1 nA for mechanism m3A3. All temperatures were set to 22°C.

In 1952, Hodgkin and Huxley showed that the relation of the channel open probability P_o_ and the test voltage *V* followed the sigmoidal equation containing the Boltzmann factor (Hodgkin & Huxley, 1952) which, through later analyses, led to the formula P_o_ = 1/(1+exp(-*q*(*V*-*V*_1/2_)/*k*_*B*_*T*)). This equation is commonly referred to as “Boltzmann equation”. Here *q* represents the electric charge, *V*_1/2_ is the half-activation potential, *k*_*B*_ is Boltzmann’s constant and *T* is the absolute temperature. This equation captures the essential features of the two-state ion permeation processes. It describes not only the channel open probability, but also the normalized current (I/I_max_) as a function of the applied voltage, which has been quite successful in fitting the patch-clamp data obtained from many ion-permeation processes and are still widely used nowadays.

Being a two-state model, the Boltzmann equation has limitations, that it does not describe the multistage or the two-direction permeation data. To use the model, all data are converted to within the range of 0 to 1 (normally done by dividing the maximum current value). However, many data also show the negative current in addition to the positive current (curves in the first three rows of Fig. 1), and some curvatures (Fig. 1) are hardly fitted by the Boltzmann equation. Therefore, more sophisticated models are developed, helpful to solve the problems in various aspects (Bezanilla, 2018; Chowdhury & Chanda, 2011, 2012; Chowdhury, Haehnel, & Chanda, 2014; Horng, Eisenberg, Liu, & Bezanilla, 2019; Islas & Sigworth, 2001; Sigg, 2014). Lacroix et al. developed a three-state model (Lacroix et al., 2012), successfully fitting the multistage charge-voltage curves obtained from many gating-current experiments of the Shaker K^+^ channel (Carvalho-de-Souza & Bezanilla, 2018; Lacroix, Hyde, Campos, & Bezanilla, 2014; Lacroix et al., 2012). Bezanilla et al. employed the sequential Boltzmann equations (two Boltzmann equations of different parameters added together) that fitted the gating-charge data of K^+^ ion permeating through the mutant Shaker channel (Bezanilla, Perozo, & Stefani, 1994). More often, the higher-rank models (usually the Markov models) with more than three-states are employed that include all possible ion-permeation pathways (Horn & Vandenberg, 1984; Vandenberg & Bezanilla, 1991; Zagotta, Hoshi, & Aldrich, 1994; Zagotta, Hoshi, Dittman, & Aldrich, 1994). These Markov models are very helpful to elucidate the allosteric ion permeation mechanisms. Although useful, these models are usually complicated containing multiple steps, and different permeation processes may employ different models, making the predicted parameters unsuitable for comparison among channels.

Therefore, a universal model is needed, not only to fit the data but also to explain the ion-permeation mechanism and compare the functions of different channels. Here we develop a generalized kinetic model. When employing three states, it is able to fit the commonly occurred current-voltage curves (typical curves are shown in Fig. 1) obtained from the patch-clamp experiments, and explain their mechanisms. The model is especially helpful to study the two-direction permeation data, which are usually left unfitted. We call this model a generalized model, because it can include other models that are commonly used nowadays. For example, when employing two states, it includes the Boltzmann equation; when employing three states, it includes the existing three-state model. In addition, this model can include the individual mechanisms suitable for each experimental design. With it, we can easily compare the functions of the different channels or the same channel under the different conditions simply by comparing the model parameters. We demonstrate the applicability of this model using several published patch-clamp data.

## Theory and Results

Transmembrane ion channels usually contain the gating and the selectivity filter domains, and some contain the extra domains sensing the change of the agonist concentrations or the environmental stimuli, such as the membrane voltage, pH, temperature, pressure, etc. Many physiological studies find that the channel can remain in the closed state (C) when its gate is closed that prohibits ion permeation, or enter the open state (O) when the gate is open that allows the binding and transmission of ions. Therefore, the open and closed states are often used in the two-state model: C↔O, as was used in deriving the Boltzmann equation (Hodgkin & Huxley, 1952).

However, ion permeation through channels can involve more than two states. Indeed, many studies find that the channel can enter the inactivation state as well. For example, the voltage-gated K^+^ channel can enter the inactivation state during the sustained depolarization stimulus (Choi, Aldrich, & Yellen, 1991; Hoshi, Zagotta, & Aldrich, 1991). The structural studies even revealed the distinct features of the ion-channel complex presumably representing the slow inactivation state, e.g., the collapsed conformation of the selectivity filter of the KcsA potassium channel (Cuello, Jogini, Cortes, & Perozo, 2010; Y. F. Zhou, Morais-Cabral, Kaufman, & MacKinnon, 2001), the lack of the ion occupation at the S1 binding site in the selectivity filter of the Kv1.2 potassium channel (Pau, Zhou, Ramu, Xu, & Lu, 2017) with an overall structure perturbed less even in the lipid bilayer environment (Matthies et al., 2018), etc. And many studies suggest that the selectivity filter can be the second gate that modulates the channel inactivation (Cordero-Morales et al., 2006; Liu, Jurman, & Yellen, 1996). Therefore, the inactivation state (I) is commonly employed in describing the ion-permeation mechanisms in three-state models, which can be expressed in several ways: C→O→I,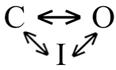, etc. However, in addition to the inactivation state, the three-state model may also employ the intermediate ion-binding state (O_i_) when the ion-binding process contains multiple steps or via the multiple subunits (such as when studying the gating charge permeating through the four voltage-sensing subunits of a voltage-gated ion channel) (Lacroix et al., 2014; Lacroix et al., 2012). In these cases, the three-state model can be expressed as C→O_i_→O or some other similar formulas. Hence the three-state model contains certainly more than one mechanism, which seems to complicate the model building procedure. Here we try to find a universal model that encompasses all these mechanisms, so that one working equation is enough to handle all types of the ion-permeation data. Aimed at deriving a universal model, we no longer use the symbols C, I, O_i_, and O in our analyses, instead, we use the general symbols like those often used in the kinetic models describing an enzymatic reaction, and denote only the ion-unbound states (E and F representing the different states of an apo channel, see captions of Figs. 2-3 for descriptions) and the ion-bound states (ES, FT, and EST representing the channels of state E or F binding ions S or T, see captions of Figs. 2-3 for descriptions) in our model. This is because any of these states can have multiple meanings suitable for the specific situation, e.g., the ES state can represent the active channel state O, or the intermediate ion-binding state O_i_, or the inactivation state I, depending on the different situations. This enables us to write the general equation for the three-state model.

**Figure 2.**
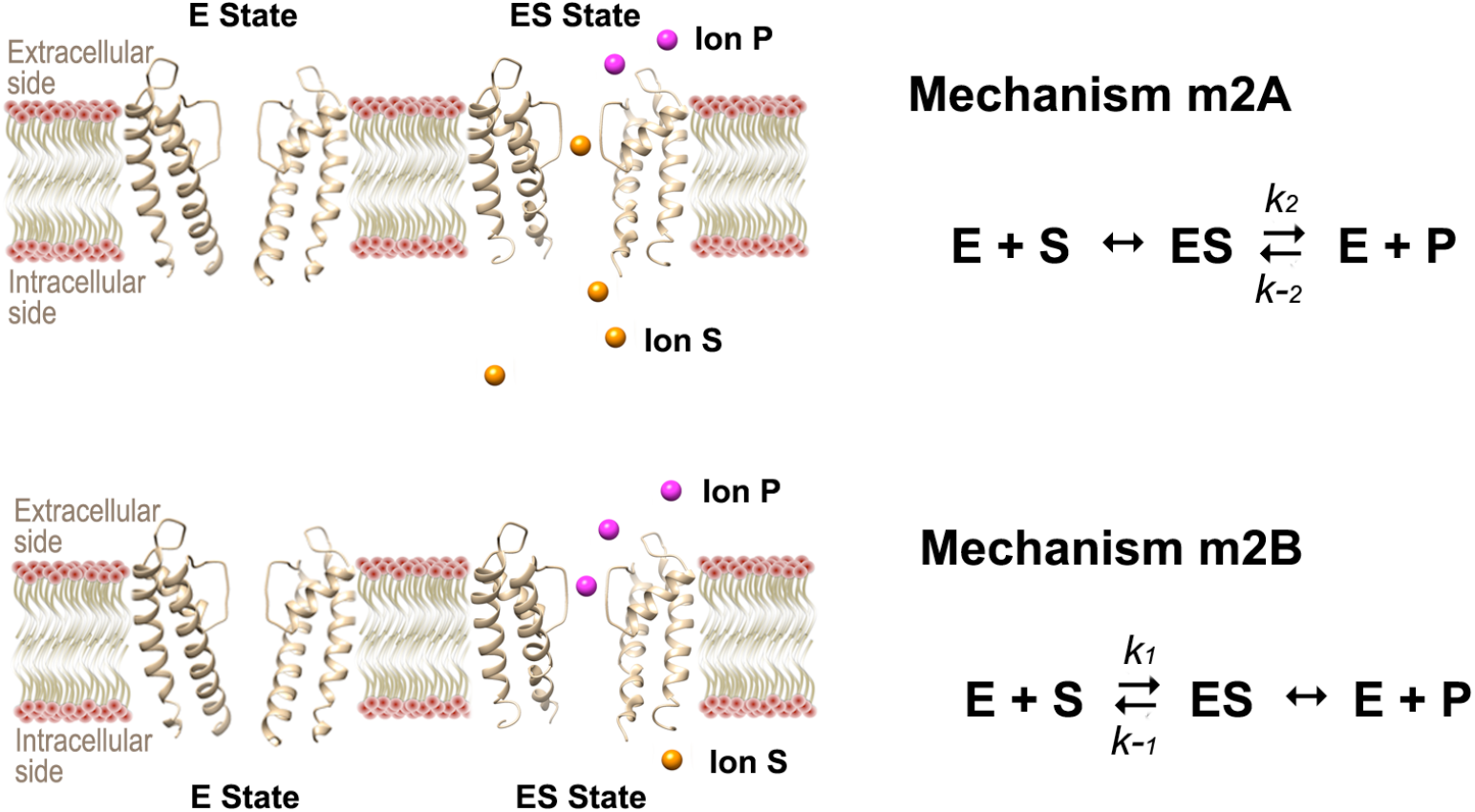
Mechanisms of the two-state model. The schematic pictures show the E- and ES-state channels embedded in the lipid bilayer. The apo channel is denoted as E, and the ion-bound form is denoted as ES. Note that the E state can also have ions bound on it (see text). The crystal structures of the KcsA K^+^ channel were used to represent the E (PDB 3FB6) and ES (PDB 3FB7) states. The substrate ion S, located at the intracellular side, changes the symbol to P (the product ion) after transmitted to the extracellular side and vice versa. The enzymatic reactions are written beside each schematic picture, describing each process involving two steps: the ion-binding and the subsequent ion-permeation step. The ion-binding step is at equilibrium whenever a double-direction arrow (↔) shows up. The outward rate constant is written above the right arrow and the inward rate constant is written below the left arrow for each ion-permeation step.

**Figure 3.**
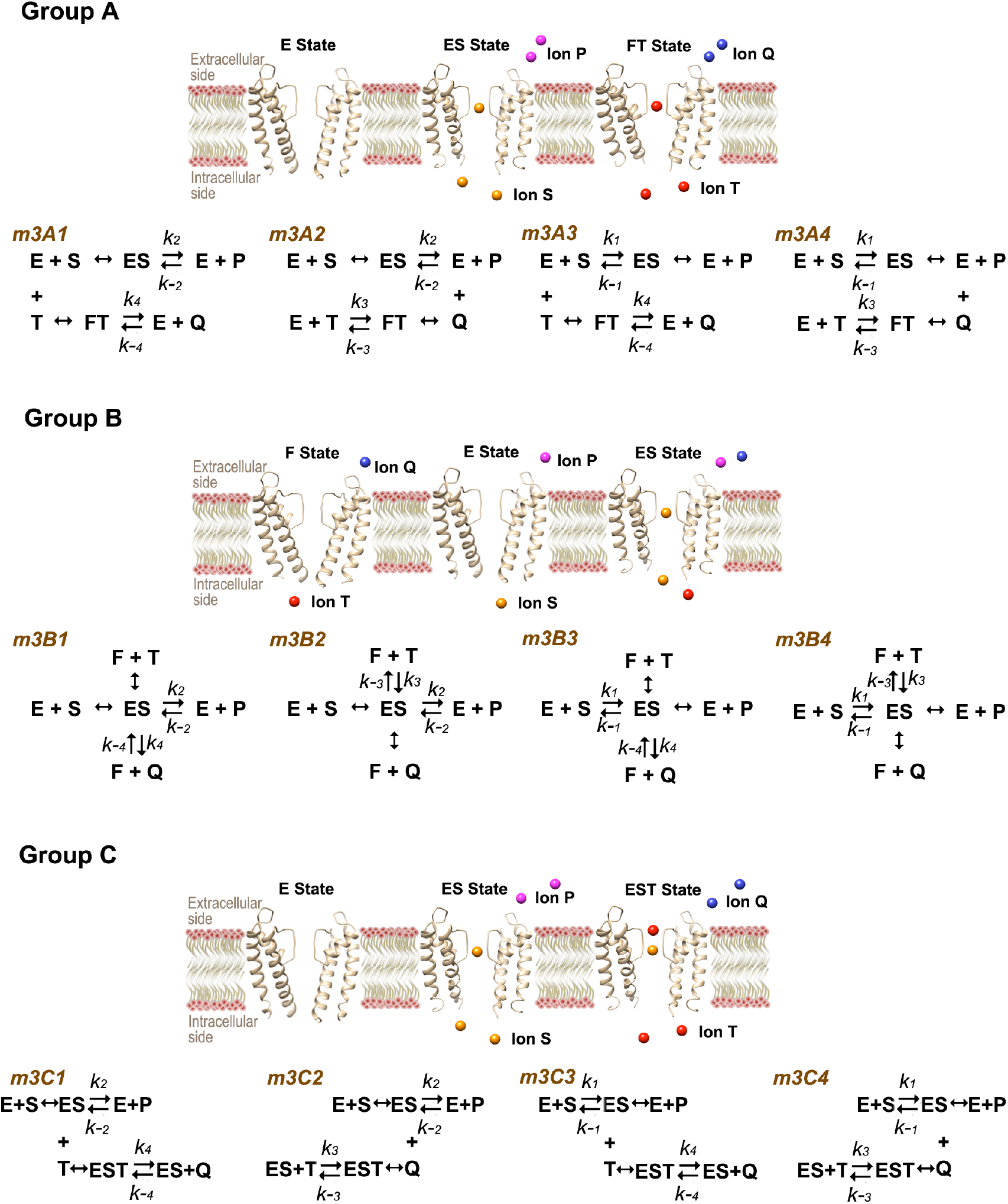
Three groups of mechanisms selected for the three-state model. Mechanisms in group A employ one ion-unbound channel state E and two ion-bound channel states ES and FT, represented by the crystal structures of KcsA K^+^ channel (PDB: 3FB6, 3FB7, and 3FB8). T can be the same or the different type of ion as S. T is also located at the intracellular side, and changes the symbol to Q after transmitted to the extracellular side. For clarity, we always use the different symbols S and T to describe the mechanism, although they may represent the same type of ion in reality. The ion-bound state (ES or FT) can have several meanings, representing such as the active channel state, the partial-active or the intermediate state, the active state with the enhanced conductivity, the inactivation state, etc. But these representations are not shown in the schematic picture. Below the picture, four mechanisms of group A are listed, each composed of a pair of equations describing the ion-binding and the subsequent ion-permeation processes. The equilibrium binding step and the relevant rate constants are described similarly as those in Fig. 2. The four mechanisms differ in the ion-permeation directions. Mechanisms in group B employ two ion-unbound states E and F (PDB 3FB5), and one ion-bound state ES, and mechanisms in group C employ one ion-unbound state E and two ion-bound states ES and EST (PDB 3FB7), where the EST state has two ions S and T bound on it. The associated mechanisms are written blow each picture. Note that the F and EST states can also have several meanings, some overlapping those represented by the ES or FT state, but we do not show their multiple meanings in this figure.

The next question is whether the three-state model is enough to handle all types of the patch-clamp data? This depends on the experimental design and on the shapes of the current-voltage curves obtained. In the normal patch-clamp experiments, the currents are recorded responding to the specified voltage elicited. These recordings do not distinguish the transition of states not involving the change of currents, such as the transitions among several closed states from C_1_→C_2_→C_3_, etc. Unless the special experimental designs are employed that can incur current changes from these states, they have to be combined into one closed state due to the lack of information. Similarly, if the transitions among several inactivation states do not incur current changes, they need be combined as well. Although the model can contain multiple open states, whether to employ all these states depends on the shape of the current-voltage curve. We suggest starting the fitting procedure with the lower-rank model whenever possible unless otherwise required by the specific experimental design. This is because several parallel or sequential steps can be combined into one step if they do not incur the appreciable current changes. For example, if the events of ion-permeation through different subunits (representing the different states) occur simultaneously that each individual state cannot be differentiated by the current curve, these states can be combined as one state. The sequential ion-binding processes E + S → ES and ES + S → ES_2_ are readily combined into one ion-binding process E + 2S → ES_2_ when the data obtained is not sufficient to differentiate the ES state.

Following these ideas, we find that the three-state model is enough to describe the commonly occurred current-voltage curves shown in Fig. 1. Therefore, we focus on explaining the two- and three-state models in this paper.

### A. The two-state model

Two mechanisms exist because the channel can conduct ions in two directions, albeit with different abilities. Here we follow the Michaelis-Menten mechanism and use the equilibrium approximation analysis. In the first mechanism (m2A in Fig. 2), the ion binding at the intracellular side is in equilibrium relative to the outward conduction step, so that *k*_2_ << *k*_-1_ and *k*_-2_[P] << *k*_1_[S]. And in the second mechanism (m2B in Fig. 2) the ion binding at the extracellular side is in equilibrium relative to the inward conduction step (*k*_2_ >> *k*_*-*1_ and *k*_-2_[P] >> *k*_1_[S]). Here we use brackets to denote the concentrations of each species, e.g., [S] and [P] represent the concentrations of the substrate ion and the product ion at the intracellular and the extracellular side, respectively. *k*_i_ is the rate constant associated with each ion-permeation step labeled on each reaction shown in Fig. 2. Note that the reaction rate is simply the current I, thus I = *k*_2_[ES] – *k*_-2_[E][P] and I = – (*k*_-1_[ES] – *k*_1_[E][S]) for the first and second mechanism, respectively. Now, let Et = [E] + [ES], we obtain a general equation for the two-state model:

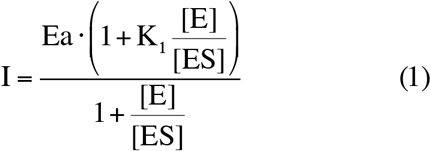

In the first mechanism, Ea = *k*_2_·Et and K_1_ = –*k*_-2_·[P]/*k*_2_. And in the second mechanism, Ea = –*k*_-1_·Et and K_1_ = –*k*_1_·[S]/*k*_-1_. Note that the expressions of Ea and K_1_ need not be identified at the stage of data fitting that each of them can be considered simply as one parameter of Eq. (1). After fitting the curve, their expressions can be used to explain and compare the channel functions.

The next step is to relate the concentration ratio [E]/[ES] to the test voltage *V*. Hodgkin and Huxley used the open probability theory (Hodgkin & Huxley, 1952), here we use the principles of chemical potentials that enable us to express all the relevant concentration ratios of the different states, such as [ES]/[E], [FT]/[E], [EST]/[ES], etc. According to the thermodynamic principles, at equilibrium, the chemical potentials of the E, S, and ES in the first step of mechanism m2A have the relation: *μ*_E_ + *μ*_S_ = *μ*_ES_. Now express each *μ* using the thermodynamic principle:

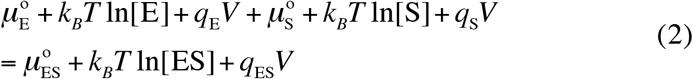

Here *μ*°_i_ denotes the chemical potential of each species *i* under the standard state, *V* is the electrostatic potential (it is the membrane potential in the ion-channel studies), *q*_*i*_ is the electric charge of each species *i, k*_*B*_ is Boltzmann’s constant, and *T* is the absolute temperature. Rearranging this equation, we obtain:

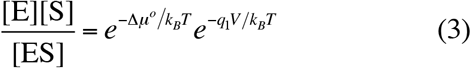

Here *q*_1_ = *q*_E_ + *q*_S_ – *q*_ES_, which represents the change of the electric charge before and after the ion binding to the protein.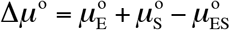. Define that [E] = [ES] when *V* = *E*_1_ (*E*_1_ is the half-activation potential, conventionally denoted as *V*_1/2_, representing the voltage at which half channels are in the E state and the other half are in the ES state), and Δ*μ*° can be expressed as a function of 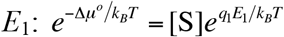. Inserting this into Eq. (3), and assuming that the solutions contain the abundant permeant ions (so that the ion concentrations change little at *V* and *E*_1_), we obtain:

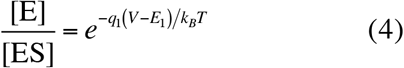

Eq. (4) is analogous to the results obtained by the probability theory (Hodgkin & Huxley, 1952), except that *q*_1_ has a slightly different meaning than the one shown in the original Boltzmann equation, here it describes the change of the charges of the ion-protein complex. Because proteins contain the charges, dipoles, quadruples, etc., and their values may also change during the ion-binding process, so the value of *q*_1_ need not be integers any more, which is often found in real cases. Normally, a large *q*_1_ value can be interpreted as a large number of ions binding to the channel. With this definition of *q*_1_, the “ion-unbound” state can still have ions bound inside the channel, and this was confirmed by many structural studies that the closed state channels bound ions inside their selectivity filters (Doyle et al., 1998; Hite et al., 2015; Tao, Avalos, Chen, & MacKinnon, 2009). Here *q*_1_ shows only the electric charges altered in the ion-protein complex in the “ion-bound” state relative to that in the “ion-unbound” state. And this is why sometimes the calculated *q*_1_ value is small that is hardly explained by the charges carried by ions. Here, the “ion-unbound” state is merely a name representing one channel state, which may not be the ion-depleted state in reality. However, for the sake of clarity, we still use the ion-depleted structure to denote the “ion-unbound” state in Figs. 2 and 3.

Performing the similar analysis for the mechanism m2B, we obtain the general formula for the two-state model that relates the current I to the test voltage *V*:

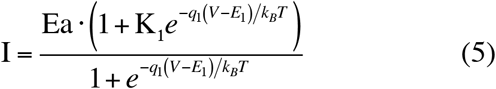

When using the normalized current and when K_1_ = 0, Eq. (5) reduces to the Boltzmann equation. Apparently, the Boltzmann equation is only a limiting case of the two-state model presented by Eq. (5), which occurs when the channel conducts ions in only one direction.

### B. The three-state model

Many mechanisms exist for the three-state model. We select three groups of mechanisms (Fig. 3) that are likely employed in real problems because they can describe all curves shown in Fig. 1. We find that the mechanisms in the first group are especially useful where the channel has one ion-unbound state E and two ion-bound states ES and FT (see Fig. 3). Here ES and FT can represent the channel having the different abilities to conduct ions in the inward and outward directions that results in rectifications (Fig. 1 B1, B2, C1-C3); or they can represent the channel having the different abilities to conduct ions in one direction that results in multistage permeations (Fig. 1 E1-E3); or one of them can represent the inactivation state (bound with ions but not transmitting them) that results in the conditional inactivation curves identified by a noticeable bell shape (Fig. 1 D1-D3). Mechanisms in the second group contain two ion-unbound states E and F, and one ion-bound state ES. Here E and F can represent the different rest-state channels having the different propensities to bind ions competing for the same conductive conformation of the channel, which can include the multistage permeation process; or one of them can represent the inactivation state that results in the conditional inactivation curves. Mechanisms in the third group contain one ion-unbound state E and two ion-bound states ES and EST, where the EST state represents the channel binding ions S and T. Mechanisms in the third group are most suitable for describing the multistage permeation process, but they are not limited to this case, e.g., ES or EST can also represent the inactivation state that results in the bell-shaped inactivation curves. Therefore, multiple mechanisms can lead to the same current-voltage curve. We may select the one that reasonably explains the ion permeation data consistent with the experimental design meanwhile yielding the smallest error in fitting the curve.

Although the mechanisms vary from one another, they all reduce to the same working equation if using the universal parameters Ea, K_1_, and K_2_ (their expressions for the individual mechanisms are listed in Appendix A):

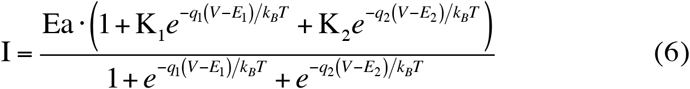

Here *q*_1_, *q*_2_, *E*_1_, and *E*_2_ have similar meanings as those described for the two-state model (see Appendix A for the definitions of *q*_1_ and *q*_2_ for each mechanism). Ea, K_1_, and K_2_ have different expressions for each mechanism (Appendix A), but they need not be identified at the stage of the data fitting. This means that we can fit the data simply using Eq. (6), then find the most suitable mechanism based on the parameters. Alternatively, we can choose several mechanisms and fit the data using their specific equations determined by the expressions of Ea, K_1_, and K_2_ (Appendix A), then select the mechanism that yields the smallest error in fitting the data. Note that when comparing the conduction rates described by the same mechanism, we can directly use the values of Ea·K_1_ and Ea·K_2_. For example, in the mechanism m3A2, Ea·K_1_ = *k*_2_·Et and Ea·K_2_ = –*k*_-3_·Et, thus comparing the absolute values of Ea·K_1_ and Ea·K_2_ is similar to comparing *k*_2_ and *k*_-3_ if Et is kept constant.

Eq. (6) is a universal working equation of the three-state model for fitting the data. Can there be a universal mechanism of the three-state model for explaining the data? Among the twelve mechanisms we have proposed (Fig. 3), we find that mechanisms m3A2 and m3A3, differing in the permeation directions, are especially useful that they describe most of the current-voltage curves appearing at different circumstances. Indeed, all curves in Fig. 1 were drawn using these two mechanisms. These curves vary from the simplest ohmic behavior (Fig. 1 A1), to the inward and outward rectifications (Fig. 1 B1, B2, C1-C3), inactivation (Fig. 1 D1-D3), and even the multistage permeations (Fig. 1 E1-E3). Employing the same mechanism enables us to compare the model parameters directly that immediately explains the altered channel functions. For example, the outward rectification curves C1 and C2 look different (Fig. 1), but they differ only in the half-activation potentials (*E*_1_ = 50 mV, *E*_2_ = -150 mV in C1 curve, and *E*_1_ = 100 mV, *E*_2_ = -200 mV in C2 curve). They look different simply because the voltage is restricted to below 100 mV. Besides the shift in the half-activation potentials, changes in *q* may also contribute to the rectification behavior within a restricted voltage range, e.g., the curve C2 itself is due to a larger number of ions binding to the channel at the intracellular side relative to that at the extracellular side (*q*_1_ = -1.5 e, *q*_2_ = 0.3 e), accompanied by the shift in the half-activation potentials (*E*_1_ = 100 mV, *E*_2_ = -200 mV). In addition, a large difference in K_1_ and K_2_ can also result in rectifications, e.g., the inward rectification curve B2 is due to a larger inward conduction rate relative to the outward conduction rate (K_2_ = - 12, K_1_ = 1). Thus, comparing model parameters enables us to compare the channel functions directly.

Note that Eq. (6) is not limited to describing the mechanisms shown in Fig. 3, it can also describe other mechanisms employing three channel states, including the previously published three-state model (Lacroix et al., 2012) (see Appendix B). Therefore, Eq. (6) is a more general form of the three-state model.

### C. The generalized kinetic model (with *N* states)

If the current curves show apparently two or more intermediate states, then the higher rank model should be used. Generally, the multi-state kinetic model can be expressed as:

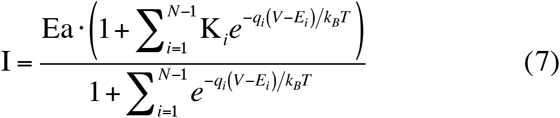

Here the parameters *q*_i_ and *E*_i_ are the electric charge changed (relative to the reference state) and the half-activation potential associated with each state *i* relative to the reference state defined for each mechanism (e.g., E state is the reference state in mechanism m3A2), and their meanings are explained analogously to those of the two- and three-state models. Like the two- and three-state models, the *N*-state model contains multiple mechanisms, and the values of Ea and K_i_ depend on the individual mechanisms.

When employing four states, Eq. (7) can include the sequential Boltzmann equations (adding up two Boltzmann equations of the different parameters) (Bezanilla et al., 1994), and their relations are presented in Appendix C. Note that in some circumstances, the four channel states can be regrouped into three states that also reasonably explains the ion-permeation data (Appendix C). Indeed, Bezanilla et al. found that the curves fitted by the sequential Boltzmann equations and the three-state model were indistinguishable in their case (Bezanilla et al., 1994), confirming that two or more states can be combined into one state if not incurring the appreciable current changes.

## Discussion

How does this model work in reality? Here we show several examples using the published patch-clamp data. The fitted curves are shown in Fig. 4 and the model parameters are shown in Table 1.

**Table 1.** Three-state model parameters selected for the current-voltage curves shown in Fig. 4.

| Curves<sup>a</sup><br>in Fig. | $q_1$ (e) | $q_2$ (e) | $E_1$ (mV) | $E_2$ (mV) | $K_1$<sup>b</sup> | $K_2$<sup>b</sup> | $Ea$<sup>c</sup> | $k'_1$<sup>d</sup> | $k'_2$<sup>e</sup> | σ<sup>f</sup> | Ref.<sup>g</sup> |
| --- | --- | --- | --- | --- | --- | --- | --- | --- | --- | --- | --- |
| 4A1 | -1.32 | 0.25 | 135 | -143 | 2.802 | -3.994 | 903 pA | 2530.206 | 3606.582 | 0.00868 | <sup>1</sup> |
| 4A2 | -1.9 | 0.25 | 120 | -165 | 1.684 | -4.996 | 1128 pA | 1899.552 | 5635.488 | 0.00675 | <sup>1</sup> |
| 4B1* | 2.227 | -0.349 | -90.652 | 81.927 | 1.228 | -4.116 | -0.044 nA | 0.054 | 0.181 | 0.00902 | <sup>2</sup> |
| 4B2* | 2.03 | -0.66 | -128.719 | 78 | 7.682 | -11.364 | -0.0408 nA | 0.313 | 0.464 | 0.0132 | <sup>2</sup> |
| 4C | -1.25 | 0.67 | 46 | -154.597 | 11.494 | -102.241 | 0.0666 μA | 0.766 | 6.809 | 0.00628 | <sup>3</sup> |
| 4D | -1.361 | 0.211 | 125.366 | -192.844 | 16 | -4.795 | 132.217 pA | 2115.472 | 633.981 | 0.00532 | <sup>4</sup> |
| 4E1* | 4.74 | -0.88 | 0 | 30 | 0 | -0.46 | -102.422 pA/pF | 0 | 47.114 | 0.0138 | <sup>5</sup> |
| 4E2* | 5.11 | -1.68 | -9.562 | 21 | 0.00466 | -0.0879 | -113.74 pA/pF | 0.53 | 9.998 | 0.0144 | <sup>5</sup> |
| 4F | -1.355 | 0.26 | -2.41 | -246.583 | 0.0222 | -14.901 | 1.799 nA | 0.0399 | 26.807 | 0.00271 | <sup>6</sup> |

<sup>a</sup> Mechanism m3A3 was employed for the curves labeled with an asterisk, and mechanism m3A2 was used for all other curves.
<sup>b</sup> $K_1$ and $K_2$ are dimensionless.
<sup>c</sup> $Ea$ has the same unit as the current reported in the source data file, which is also written following each $Ea$ value in the table.
<sup>d</sup> $k'_1 = Et \cdot k_2 = |Ea \cdot K_1|$ for mechanism m3A2, and $k'_1 = Et \cdot k_{-1} = |Ea \cdot K_1|$ for mechanism m3A3. $k'_1$ has the same unit as $Ea$ .
<sup>e</sup> $k'_2 = Et \cdot k_{-3} = |Ea \cdot K_2|$ for mechanism m3A2, and $k'_2 = Et \cdot k_4 = |Ea \cdot K_2|$ for mechanism m3A3. $k'_2$ has the same unit as $Ea$ .
<sup>f</sup> See Materials and Methods for the definition of $\sigma$ .
<sup>g</sup> Ref. 1 (Lemoine et al., 2020); Ref. 2 (Zheng et al., 2020); Ref. 3 (Chiasson et al., 2017); Ref. 4 (Syrjanen et al., 2020); Ref. 5 (Huang et al., 2019); Ref. 6 (Zhou et al., 2017).

**Figure 4.**
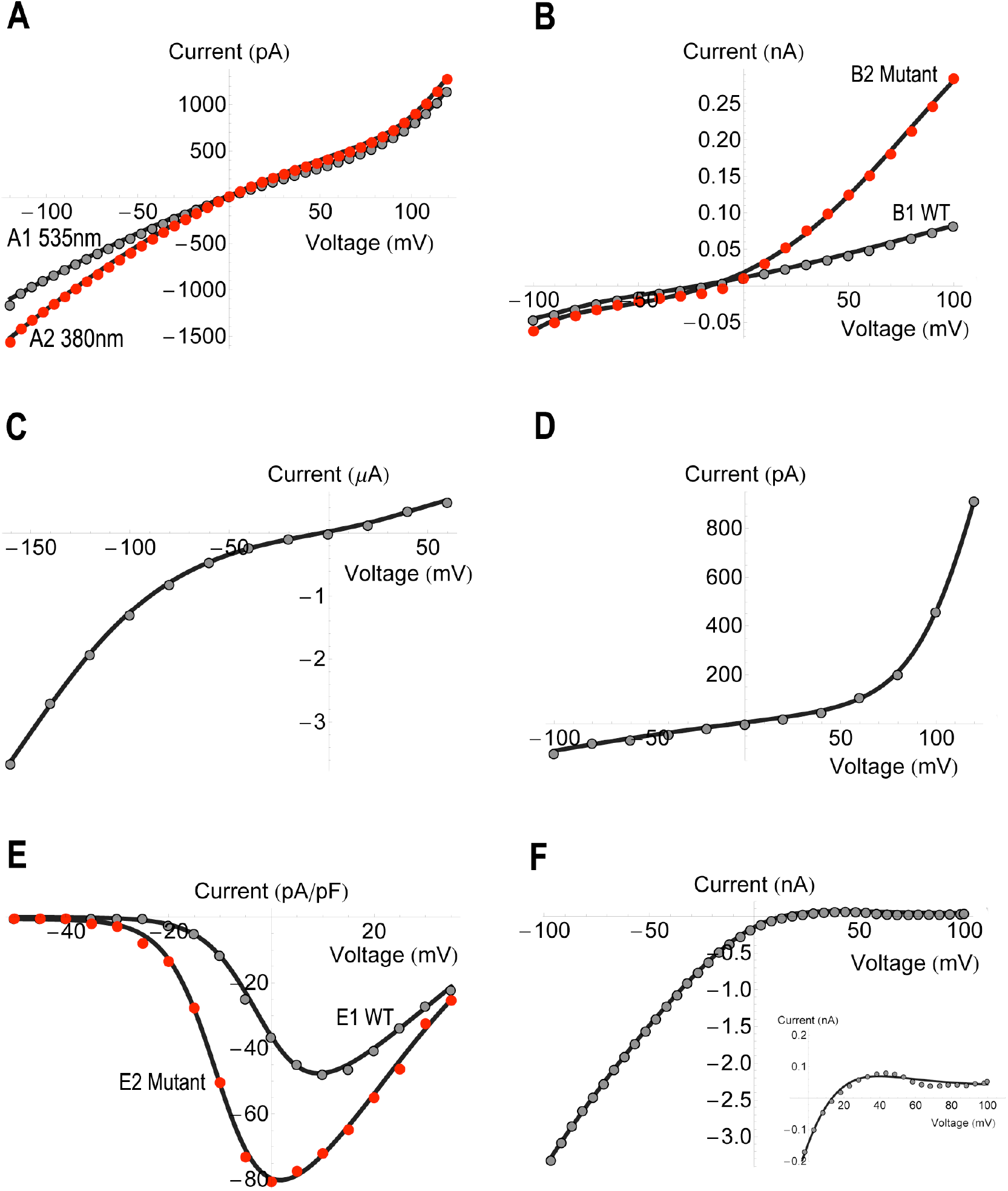
Experimental current-voltage curves fitted by the three-state model. Source data are represented by the filled circles, and the model data are plotted by the solid black curves. Reference to the source data, the selected mechanism, and the fitted parameters are listed in Table 1. The room temperature (22°C) was used for all curve-fitting procedures. The inset in Panel F shows the enlarged plot of a local region (positive voltages) of the current-voltage curve.

### Two-direction permeation curves

Lemoine et al. have designed an optical molecule attached close to the pore of the GluD2 receptor that senses the light of the specific wavelength to occlude or open the entrance to the channel pore (Lemoine et al., 2020). The two current curves under the 535 nm and 380 nm light illuminations (source data from Figure 4A of Ref. (Lemoine et al., 2020)) were fitted with the three-state model and presented in Fig. 4A. Mechanism m3A2 is suitable to explain these curves. It describes the two-direction ion permeations via the two ion-bound states ES and FT. Here ES represents the channel state inclined to direct the outward current, and FT represents the channel state inclined to direct the inward current. Compared with the 535 nm light illumination, the 380 nm light increases the inward conduction rate (*k*′_2, 535nm_ = 3606.58 pA, *k*′_2, 380nm_ = 5635.49 pA, see captions of Table 1 for the definition of *k*′) and decreases the outward conduction rate (*k*′_1, 535nm_ = 2530.21 pA, *k*′_1, 380nm_ = 1899.55 pA), assuming that Et is unchanged. This produces the increased inward current at negative potentials, reflecting the optical molecule’s ability to sense the specific light and magnify the inward current. We also note that it modifies the ion binding at the intracellular side (*q*_1, 535nm_ = -1.32 e, *q*_1, 380nm_ = -1.9 e) that more ions bind to the channel under the 380 nm light illumination (when the optical molecule switches to the cis-conformation that opens the pore entrance), consistent with the lock property of the optical molecule. In addition, the guanidinium moiety of the optical molecule may also play a role to bring about these changes, possibly through interacting with the positively charged ions upon switching its locations under the different light illuminations.

Zheng et al. solved the cryo-EM structures of a eukaryotic cyclic nucleotide-gated channel TAX-4, and found that the double mutations of the hydrophobic residues F403V and V407A in the cavity of the channel can increase the outward basal current (Zheng et al., 2020). The current curves of the wild-type and mutant channels (source data from Figure 4a in Ref. (Zheng et al., 2020)) were fitted with the three-state model and plotted in Fig. 4B. Mechanism m3A3 is suitable for this case, where the ES and FT states describe the channel abilities for the inward and outward conductions, respectively. The cryo-EM structures reveal that F403 and V407 form the hydrophobic gate inside the channel cavity that they block the permeation pathway during the closed state but pave the way for ion permeation by rotating aside upon switching to the open state (Zheng et al., 2020). Mutations to valine and alanine certainly increase the central space in the cavity that makes the channel a bit “leaky” in a sense (Zheng et al., 2020). Comparing the model parameters, the mutations increase the conduction rates in both directions (*k*′_1, wt_ = 0.054 nA, *k*′_1, mutant_ = 0.31 nA, and *k*′_2, wt_ = 0.18 nA, *k*′_2, mutant_ = 0.46 nA), conforming the “leaky” property of the mutant channel. In addition, ion binding at the intracellular side is roughly doubled (*q*_2, wt_ = -0.35 e, *q*_2, mutant_ = -0.66 e), but that of the extracellular side is slightly decreased (*q*_1, wt_ = 2.23 e, *q*_1, mutant_ = 2.03 e), reflecting the increased ability of the mutant channel for the outward conduction. This, together with the increased conduction rates in both directions, exhibits the increased outward basal current as the overall effect.

### Rectification curves

Chiasson et al. have reported a *brush* mutation in the cyclic nucleotide-gated channel that resulted in the gain-of-function, manifested by the inward rectification of the Ca^2+^ current (Chiasson et al., 2017). The typical current curve (source data from the *brush* curve of Figure 3D in Ref. (Chiasson et al., 2017)) was fitted by the three-state model and plotted in Fig. 4C. Employing mechanism m3A2, the inward rectification is mainly due to the larger inward conduction rate (K_2_ = -102.24) relative to the smaller outward conduction rate (K_1_ = 11.49).

Syrjanen et al. have reported the structure of a calcium homeostasis modulator that produced the outward-rectification current (Syrjanen et al., 2020). The typical current curve (source data from the hCALHM1 curve of Figure 1b in Ref. (Syrjanen et al., 2020)) was fitted by the three-state model (mechanism m3A2) and plotted in Fig. 4D. This time, the outward rectification is due to the larger number of ions binding at the intracellular side (q_1_ = -1.36 e, q_2_ = 0.21 e), and a larger outward conduction rate (K_1_ = 16, K_2_ = -4.8).

The above results, together with the model curves shown in Fig. 1, show the varied reasons that lead to the rectifications. Theoretically, a large difference in K_1_ and K_2_ can lead to rectifications. But these rectifications may not appear within the restricted voltage range selected for the current recordings. Thus the rectification curve shown up in a narrowed voltage range is usually due to a collective action mixed with the individual changes in *q, E*, and K. Simply comparing *q*_1_ and *q*_2_ or comparing K_1_ and K_2_ sometimes can lead to inconsistent conclusions, e.g., in Fig. 4C, a larger *q*_1_ compared to *q*_2_ (q_1_ = -1.25 e, q_2_ = 0.67 e) does not lead to the conclusion of the outward rectification.

### Bell-shaped curves

Huang et al. have reported a mutation of the voltage-gated calcium channel that resulted in the gain-of-function and produced the increased inward current (Huang et al., 2019). The current curves of the wild-type and mutant channels (source data from Figure 3b of Ref. (Huang et al., 2019)) were plotted in Fig. 4E. We find that the mechanism m3A3 is suitable to explain these curves. Here ES represents the inactivation channel state at the negative potentials. Note that the “inactivation state” used here is only a general term denoting one nonconducting channel state, which can result from several conditions including the closing of the inner gate. As the test voltage increases, the channel gradually recovers from the inactivation state and switches to the E state then to the FT state, which is accompanied by the inward conduction of ions at the less negative to the small positive potentials. At the more positive potentials, the FT state dominates that directs the outward current. This is how the bell-shape curve is formed around 0 mV. Comparing the parameters of the wild-type and mutant channel curves, the mutation increases both outside and inside ion binding (*q*_1, wt_ = 4.74 e, *q*_1, mutant_ = 5.11 e; *q*_2, wt_ = -0.88 e, *q*_2, mutant_ = -1.68 e), shifts both half-activation potentials to the left (*E*_1, wt_ = 0 mV, *E*_1, mutant_ = -9.56 mV; *E*_2, wt_ = 30 mV, *E*_2, mutant_ = 21 mV), and significantly decreases the outward ion-conduction rate (*k*′_2, wt_ = 47.11 pA/pF, *k*′_2, mutant_ = 10 pA/pF). These changes lead to the larger inward current shifted to the left of the wild-type current curve (Fig. 4E), which can be interpreted as the gain-of-function.

Zhou et al. have solved the cryo-EM structures of the human endolysosomal TRPML3 channel and studied its functions (Zhou et al., 2017). The ligand-activated current curve of the wild-type channel (source data from the ML-SA1 ligand-bound current curve in Figure 2b of Ref. (Zhou et al., 2017)) showed not only an inward rectification, but also a bell curvature at positive potentials (Fig. 4F, the inset shows the enlarged plot of the current at positive potentials). We can use the mechanism m3A2 to explain this phenomenon. At negative potentials, the FT state is the active form that directs the large inward currents (K_2_ = -14.9). As the test voltage increases, the ion permeation changes the direction, and the FT state gradually switches to the E then to the ES state. The ES state is the inactivation state because the channel nearly prohibits ion conduction at the very positive potentials (K_1_ = 0.022).

### Some further points

The above studies show only the applicability of the three-state model, that it can fit the curve and explain their mechanisms in accordance with the experimental findings. But they tell nothing about the reliability of the model parameters, because only one curve (the averaged data in most cases) is used for each case in our study. The reliability of the model parameters depends on the reproducibility of each data set, and hence is beyond the scope of this paper. However, the model parameters do provide a way to examine the reproduced data sets (by comparing the model parameters obtained from fitting the curve using each data set) and check whether these data values are consistent with one another.

Many patch-clamp recordings show the continuously increasing currents not reaching the saturation level, and we may wonder when the current will saturate. This information is easily obtained from the three-state model if the parameters are reliable. For example, in mechanism m3A2, the saturation currents are Ea·K_1_ and Ea·K_2_ at the positive and negative potentials, respectively. The saturation currents can occur within or beyond the physiological recording range, and the model can help predict these values or help record these values based on the predicted voltage range.

Do the above analyses confirm that the channels switch among only three possible states in those ion-permeation processes? No, the model only suggests that each permeation event employs only three major channel states, that the currents elicited from the minor states are either too small to be detected or merged into the major states during the concerted movements. Employing more states certainly is possible, but not suggested because extra parameters can incur large inaccuracies if not supported by the enhanced recordings that yield adequate curvatures in the current-voltage curves to discriminate the intermediate states. However, these analyses do suggest a simplified but universal mechanism, sufficient to explain most ion-permeation events with only three major channel states. The essential feature of this mechanism is the voltage-dependent switch between the two ion-bound states ES and FT, reflecting channel’s altered abilities to conduct ions in the uni- or opposite directions. In fact, their presumable equivalents are already found in the structural studies. Cuello et al. have reported a series of the crystal structures of the KcsA K^+^ channel (Cuello et al., 2010), differing in the opening scale at the intracellular gate, some even accompanied by the structural change in the selectivity filter, that any of these structures may represent the ES or FT state with the altered abilities to conduct ions.

## Appendices

### Appendix A: Parameters of the three-state model

The four mechanisms in group A employ the concentration ratios [ES]/[E] and [FT]/[E]. Following similar analyses using the thermodynamic principles (as those used in deriving Eq. (4)), we obtain: 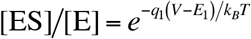 and 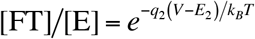. Here *q*_1_ = *q*_ES_ – *q*_E_ – *q*_S_ for mechanisms m3A1 and m3A2, and *q*_1_ = *q*_ES_ – *q*_E_ – *q*_P_ for mechanisms m3A3 and m3A4. *q*_2_ = *q*_FT_ – *q*_E_ – *q*_T_ for mechanisms m3A1 and m3A3, and *q*_2_ = *q*_FT_ – *q*_E_ – *q*_Q_ for mechanisms m3A2 and m3A4. *E*_1_ and *E*_2_ are the half-activation potentials associated with the portion of the channels changing from the E state to the ES and FT states, respectively. The expressions of Ea, K_1_, and K_2_ for the individual mechanisms of this group are:

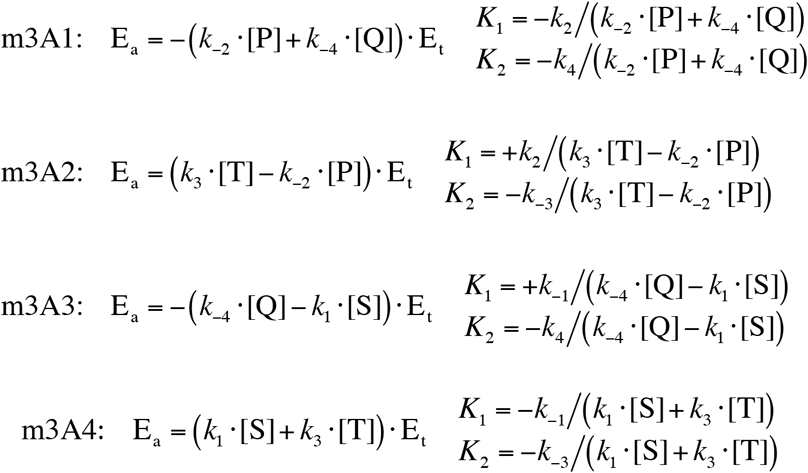

The four mechanisms in group B employ the concentration ratios [E]/[ES] and [F]/[ES]. Here 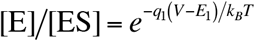 and 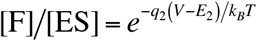, where *q*_1_ = *q*_E_ + *q*_S_ – *q*_ES_ for mechanisms m3B1 and m3B2, and *q*_1_ = *q*_E_ + *q*_P_ – *q*_ES_ for mechanisms m3B3 and m3B4. *q*_2_ = *q*_F_ + *q*_T_ – *q*_ES_ for mechanisms m3B1 and m3B3, and *q*_2_ = *q*_F_ + *q*_Q_ – *q*_ES_ for mechanisms m3B2 and m3B4. *E*_1_ and *E*_2_ are the half-activation potentials associated with the portion of the channels changing from the ES state to the E and F states, respectively. The expressions of Ea, K_1_, and K_2_ for the individual mechanisms of this group are:

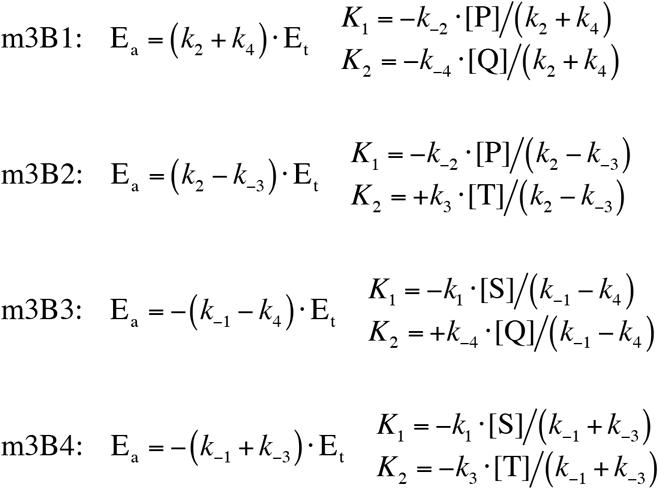

The four mechanisms in group C employ the concentration ratios [E]/[ES] and [EST]/[ES]. Here 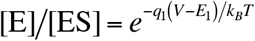 and 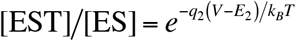, where *q*_1_ = *q*_E_ + *q*_S_ – *q*_ES_ for mechanisms m3C1 and m3C2, and *q*_1_ = *q*_E_ + *q*_P_ – *q*_ES_ for mechanisms m3C3 and m3C4. *q*_2_ = *q*_EST_ – *q*_ES_ – *q*_T_ for mechanisms m3C1 and m3C3, and *q*_2_ = *q*_EST_ – *q*_ES_ – *q*_Q_ for mechanisms m3C2 and m3C4. *E*_1_ and *E*_2_ are the half-activation potentials associated with the portion of the channels changing from the ES state to the E and EST states, respectively. The expressions of Ea, K_1_, and K_2_ for the individual mechanisms of this group are:

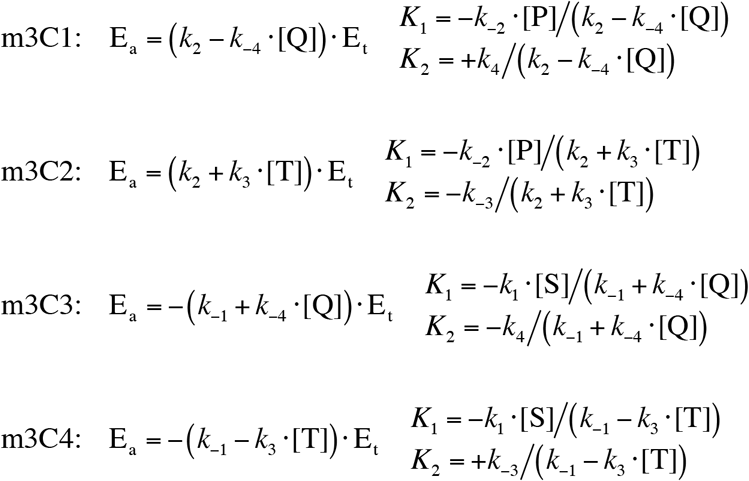

### Appendix B: Relation of the three-state models

Lacroix et al. developed a three-state model suitable to calculate the multistage gating charge as a function of the test voltage (Lacroix et al., 2012). The gating charge per voltage-sensing domain is defined as:

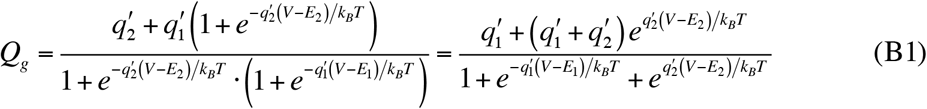

In their definitions, *q*_1_′ is the charge associated with the transition from the resting to the intermediate state, and *q*_2_′ is the charge associated with the subsequent transition from the intermediate to the active state. Now let’s define the total gating charge per voltage-sensing domain *Q*_*g*,max_ = *q*_1_′+*q*_2_′, and the fraction of the charge in the intermediate state *f* = *q*_1_′/ *Q*_*g*,max_, then we obtain:

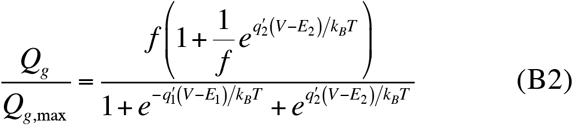

Now we can relate Eq. (B2) to mechanism m3C1:

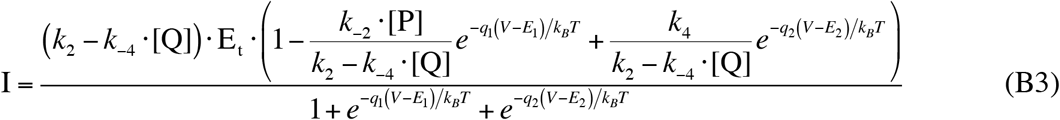

If *k*_-2_·[P] is small enough or *k*_-2_·[P] << *k*_2_ – *k*_-4_·[Q], the second term in the numerator of Eq. (B3) can be neglected, and we obtain

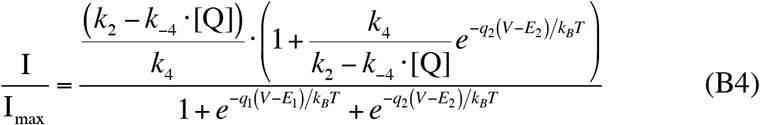

Here I_max_ = *k*_4_·Et. Comparing Eq. (B4) with Eq. (B2), *q*_1_ = *q*_1_′ and *q*_2_ = -*q*_2_′, because we have defined the transitions both starting from the intermediate state in mechanism m3C1. The fraction of charge *f* in Eq. (B2) is equivalent to the fraction of the rate constant (*k*_2_ − *k*_−4_ ·[Q]) *k*_4_ in Eq. (B4). Therefore Eq. (B2) can be considered as a special case of mechanism m3C1 defined by Eq. (B3) when *k*_-2_·[P] ≈ 0 or *k*_-2_·[P] << *k*_2_ – *k*_-4_·[Q].

### Appendix C: Relation of the four-state model and the sequential Boltzmann equations

The sequential Boltzmann equations proposed by Bezanilla et al. were also used to calculate the gating charge of the Shaker channel, that yielded a charge-voltage curve identical to that obtained by the three-state model (Bezanilla, Perozo, & Stefani, 1994). The sequential Boltzmann equations can represent the two independent ion-binding processes, that include an intermediate ES state with the faction of charge *f = q*_1_′/*Q*_*g*,max_, where *Q*_*g*,max_ = *q*_1_′ + *q*_2_′:

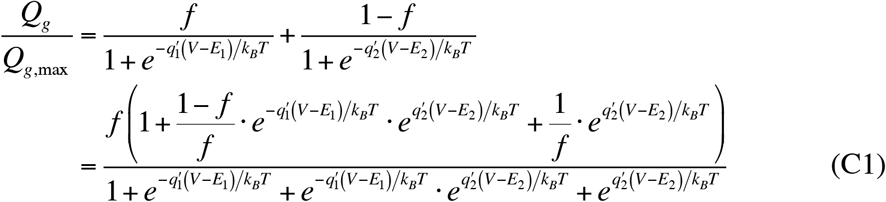

And this is equivalent to a four-state model involving the E, ES, FT, and EST states. Let’s write one simple mechanism for this four-state model:

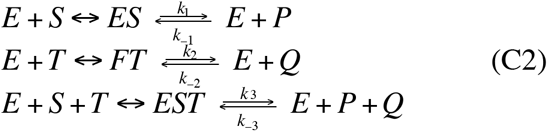

If we define I_max_ = *k*_3_·E_t_, then the normalized current for this mechanism is:

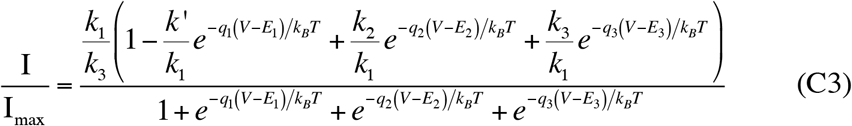

Note that Eq. (C3) is only a subset case defined by the generalized kinetic model (Eq. (7)) involving four states. Here *k*′ = *k*_-1_·[P] + *k*_-2_·[Q] + *k*_-3_·[P] ·[Q], 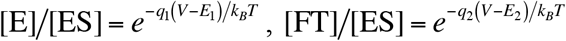, and 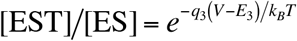. When *k*′ is small enough or *k*′ << *k*_1_, the second term in the numerator can be neglected, and we obtain:

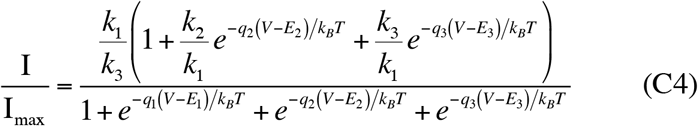

For the independent binding processes defined by Eq. (C2), 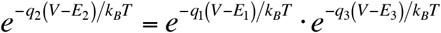, inserting this into Eq. (C4), we obtain:

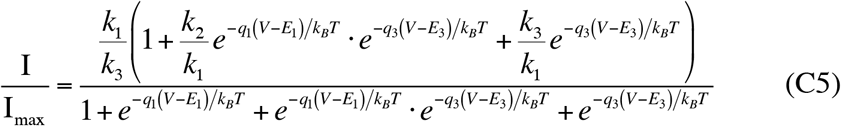

Now comparing Eq. (C5) with Eq. (C1), *q*_1_ = *q*_1_′, *q*_3_ = -*q*_2_′. The fraction of charge *f* at the intermediate state is equivalent to the rate constant ratio *k*_1_/*k*_3_, and (1-*f*)/*f* is equivalent to (*k*_2_/*k*_3_)/(*k*_1_/*k*_3_) = *k*_2_/*k*_1_. So the sequential Boltzmann equations can be described by the four-state model defined by Eq. (C5), which is a special case of Eq. (C3) that occurs when *k*′ ≈ 0 or *k*′ << *k*_1_. Therefore the sequential Boltzmann equations can be included in the general kinetic model defined by Eq. (7).

Now let’s compare Eq. (C1) and Eq. (B2). In Eq. (C1), if the second term 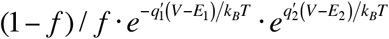 is much smaller than the other terms in the numerator, and the third term 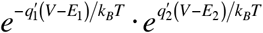 is much smaller than the other terms in the denominator, they can be neglected from the numerator and denominator, and hence Eq. (C1) is reduced to Eq. (B2). This is when the three-state model and the sequential Boltzmann equations yield the indistinguishable charge-voltage curves like those found by Bezanilla et al. (Bezanilla, Perozo, & Stefani, 1994). Following the mechanism defined by Eq. (C2), this situation can happen when the FT state is merged into another ion-bound state, so that the four-state model is readily reduced to a three-state model involving only the E, ES, and EST states. And this is why we suggest using the lower-rank model whenever the extra intermediate states cannot be differentiated by the current-voltage curves.

## Materials and Methods

Source data of the current-voltage curves plotted in Fig. 4 were obtained from the publications listed in Table 1, which were also cited in the Discussion section. For most of them, we directly used the source data as presented in the source data file along with the publications, except for the following data.

The source data of the current-voltage curves in Figure 4A of Ref. (Lemoine et al., 2020) each contained 400 pair of values. We had reduced each data size to 41 and used them for the curve fitting procedure. The data of the reduced size are presented in the supplementary table.

The source data of Figure 2b in Ref. (Zhou et al., 2017) contained 395 pair of values, and we had reduced the data size to 41. The data with the reduced size are presented in the supplementary table.

All source data were fitted following the method of the nonlinear least squares. The model parameters that yielded the smallest σ value were selected, where σ is defined as:

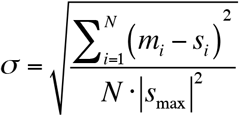

Here *s*_*i*_ represents the *i*-th current value in the source data of size *N. m*_*i*_ represents the *i*-th current value calculated by the selected three-state model. |*s*_max_| is the absolute value of the maximum current in the source data file. The parameters selected for each curve, together with the calculated σ value, are presented in Table 1.

## Acknowledgments

I am grateful to the authors of the publications listed in Table 1, who generously published their source data and shared them with the society, that made the proposed model testable.

This work was supported by Natural Science Foundation of China (Grant No. 21003023).

