## Supplementary Table for "A generalized kinetic model describes ion-permeation mechanisms in various ion channels"

The following table contains the modified data used for the curve fitting procedures. These data were adapted from the source data of Ref. 1 (Lemoine et al., 2020) and Ref. 6 (Zhou et al., 2017).

| Fig. 4A1 | | Fig. 4A2 | | Fig. 4F | |
|---|---|---|---|---|---|
| Original data from Ref. 1 | | Original data from Ref. 1 | | Original data from Ref. 6 | |
| Voltage (mV) | Current (pA) | Voltage (mV) | Current (pA) | Voltage (mV) | Current (nA) |
| -119.7517591 | -1147.412925 | -119.7517591 | -1545.956355 | -96.5575 | -3.30017 |
| -113.7781388 | -1020.394405 | -113.7781388 | -1403.53387 | -91.5835 | -3.0603 |
| -107.7438746 | -949.3916263 | -107.7438746 | -1309.709102 | -86.609 | -2.83569 |
| -101.762038 | -885.5444975 | -101.762038 | -1221.324092 | -81.5735 | -2.62695 |
| -95.7629864 | -825.2214276 | -95.7629864 | -1136.845341 | -76.5685 | -2.44324 |
| -89.78584485 | -766.7759102 | -89.78584485 | -1056.440681 | -71.4415 | -2.26135 |
| -83.789532 | -705.7741837 | -83.789532 | -972.6073677 | -66.5285 | -2.06726 |
| -77.79869666 | -648.9039503 | -77.79869666 | -892.9122414 | -61.493 | -1.89209 |
| -71.76873623 | -590.7083864 | -71.76873623 | -815.7806068 | -56.61 | -1.7157 |
| -65.77203213 | -538.0786431 | -65.77203213 | -737.9485756 | -51.4525 | -1.54419 |
| -59.84653571 | -483.4042242 | -59.84653571 | -659.9303871 | -46.63085 | -1.38 |
| -53.79662147 | -428.1645068 | -53.79662147 | -584.3612354 | -41.56495 | -1.20667 |
| -47.80969864 | -374.6962582 | -47.80969864 | -510.3713718 | -36.499 | -1.05225 |
| -41.80595202 | -321.9894972 | -41.80595202 | -436.1312656 | -31.70775 | -0.88989 |
| -35.80337916 | -269.7114358 | -35.80337916 | -362.5274494 | -26.5198 | -0.74524 |
| -29.8422789 | -216.8448208 | -29.8422789 | -292.4255213 | -21.5454 | -0.60669 |
| -23.80292844 | -167.9222383 | -23.80292844 | -224.174489 | -16.51 | -0.46997 |
| -17.8312644 | -116.728278 | -17.8312644 | -158.1252973 | -11.6272 | -0.36072 |
| -11.81343274 | -69.37953066 | -11.81343274 | -92.72307962 | -6.34765 | -0.25879 |
| -5.819076147 | -23.80806977 | -5.819076147 | -31.50634667 | -1.67847 | -0.16785 |
| 0.151414138 | 20.9612147 | 0.151414138 | 25.68969534 | 3.38745 | -0.09827 |
| 6.173158303 | 63.58555022 | 6.173158303 | 79.60814934 | 8.39235 | -0.04333 |
| 12.16477614 | 103.4415043 | 12.16477614 | 131.816094 | 13.30565 | -0.00671 |
| 18.165784 | 142.2714889 | 18.165784 | 180.2841093 | 18.34105 | 0.02136 |
| 24.15466309 | 179.8967543 | 24.15466309 | 225.6606922 | 23.2544 | 0.04211 |
| 30.16779973 | 215.3828797 | 30.16779973 | 267.7047588 | 28.35085 | 0.06042 |
| 36.1312475 | 251.047756 | 36.1312475 | 307.8430035 | 33.3557 | 0.0708 |
| 42.15886043 | 285.8610466 | 42.15886043 | 346.0327007 | 38.5132 | 0.07935 |
| 48.13834948 | 320.5508121 | 48.13834948 | 383.3953693 | 43.5486 | 0.0824 |
| 54.16752742 | 355.9788114 | 54.16752742 | 423.57328 | 48.4619 | 0.07874 |
| 60.12667142 | 395.9437561 | 60.12667142 | 462.5472755 | 53.3445 | 0.07019 |
| 66.10029172 | 437.3691842 | 66.10029172 | 506.9762901 | 58.4105 | 0.05493 |
| 72.11421086 | 483.3213886 | 72.11421086 | 554.5486168 | 63.4155 | 0.04578 |
| 78.12734751 | 533.3862021 | 78.12734751 | 607.569857 | 68.268 | 0.0415 |
| 84.09822904 | 594.0377059 | 84.09822904 | 669.4854454 | 73.578 | 0.04211 |
| 90.10119316 | 657.957497 | 90.10119316 | 737.6083046 | 78.2775 | 0.04517 |
| 96.08928974 | 730.9308297 | 96.08928974 | 820.440638 | 83.3435 | 0.04517 |
| 102.0887326 | 819.0801275 | 102.0887326 | 915.8019485 | 88.379 | 0.04333 |
| 108.0960005 | 917.8073861 | 108.0960005 | 1026.321355 | 93.292 | 0.04883 |
| 114.0739245 | 1034.598158 | 114.0739245 | 1156.628362 | 98.297 | 0.05066 |
| 119.4380575 | 1155.210892 | 119.4380575 | 1291.564876 | 100 | 0.05554 |
